# Synaptogyrin-3 Prevents Cocaine Addiction and Dopamine Deficits

**DOI:** 10.1101/2024.07.27.605436

**Authors:** Emily G. Peck, Katherine M. Holleran, Alyson M. Curry, Kimberly M. Holter, Paige M. Estave, Jonathon P. Sens, Jason L. Locke, Olivia A. Ortelli, Brianna E. George, Monica H. Dawes, Alyssa M. West, Nancy J. Alexander, Drew D. Kiraly, Sean P. Farris, Robert W. Gould, Brian A. McCool, Sara R. Jones

## Abstract

Synaptogyrin-3, a functionally obscure synaptic vesicle protein, interacts with vesicular monoamine and dopamine transporters, bringing together dopamine release and reuptake sites. Synaptogyrin-3 was reduced by chronic cocaine exposure in both humans and rats, and synaptogyrin-3 levels inversely correlated with motivation to take cocaine in rats. Synaptogyrin-3 overexpression in dopamine neurons reduced cocaine self-administration, decreased anxiety-like behavior, and enhanced cognitive flexibility. Overexpression also enhanced nucleus accumbens dopamine signaling and prevented cocaine-induced deficits, suggesting a putative therapeutic role for synaptogyrin-3 in cocaine use disorder.

## Main Text

While total drug overdose deaths in the United States are stabilizing after a decade of growth, cocaine-related deaths are on the rise^1^ and there are still no FDA-approved pharmacotherapies for cocaine use disorder (CUD). Together, this emphasizes an urgent need to understand the neurobiological underpinnings of CUD and identify new targets for drug development. Like other drugs of abuse, cocaine exerts its reinforcing properties through increased mesolimbic dopamine signaling, particularly in the nucleus accumbens (NAc)^2–4^. Chronic cocaine exposure induces compensatory adaptations in the dopamine system, resulting in hypodopaminergia^5,6^, which drives greater CUD symptom severity, such as escalated use and relapse^7,8^. While cocaine modulates dopamine signaling in multiple ways^9–13^, a great deal of research has focused on cocaine’s inhibition of the dopamine transporter (DAT), which terminates dopamine signaling and regulates extracellular dopamine levels^14–18^. However, prospective CUD pharmacotherapies aimed directly at the DAT often have high abuse potential^19^. Therefore, focusing on indirect modulators of mesolimbic dopamine function may yield drug targets with low abuse potential and high therapeutic value. Here, we investigate an understudied protein, synaptogyrin-3 (Syngr3), as a novel modulator of dopamine function and cocaine-taking behavior.

Syngr3 is a synaptic vesicle protein that has recently been associated with traumatic brain injury and neurodegenerative diseases^19–22^, but its exact function(s) remains obscure. While its better-known family member, synaptogyrin-1, facilitates vesicle priming and membrane fusion ubiquitously throughout the brain^23–25^, Syngr3 is highly enriched in midbrain dopamine neurons^26^, suggesting a more specialized role in dopamine transmission. Indeed, Syngr3 is uniquely positioned within the presynaptic terminal to interact with various aspects of vesicle dynamics. For instance, Syngr3 is thought to play a role in exocytosis^23,27^ and might link storage vesicles to the cytoskeleton through its binding to microtubule-associated tau^21,28^. Additionally, Syngr3 may accelerate dopamine vesicle refilling by co-localizing release and reuptake sites via its interactions with the DAT and vesicular monoamine transporter 2 (VMAT2; Fig. 1A)^29,30^. Because of its unique position in dopamine terminals, we investigated the role of Syngr3 in CUD-related behavior and dopamine neurotransmission. We found that high cocaine use was associated with low Syngr3 levels, and viral overexpression of Syngr3 in dopamine neurons boosted overall signaling and protected against hypodopaminergia, ultimately leading to mitigation of addiction-like behaviors.

**Fig. 1.**
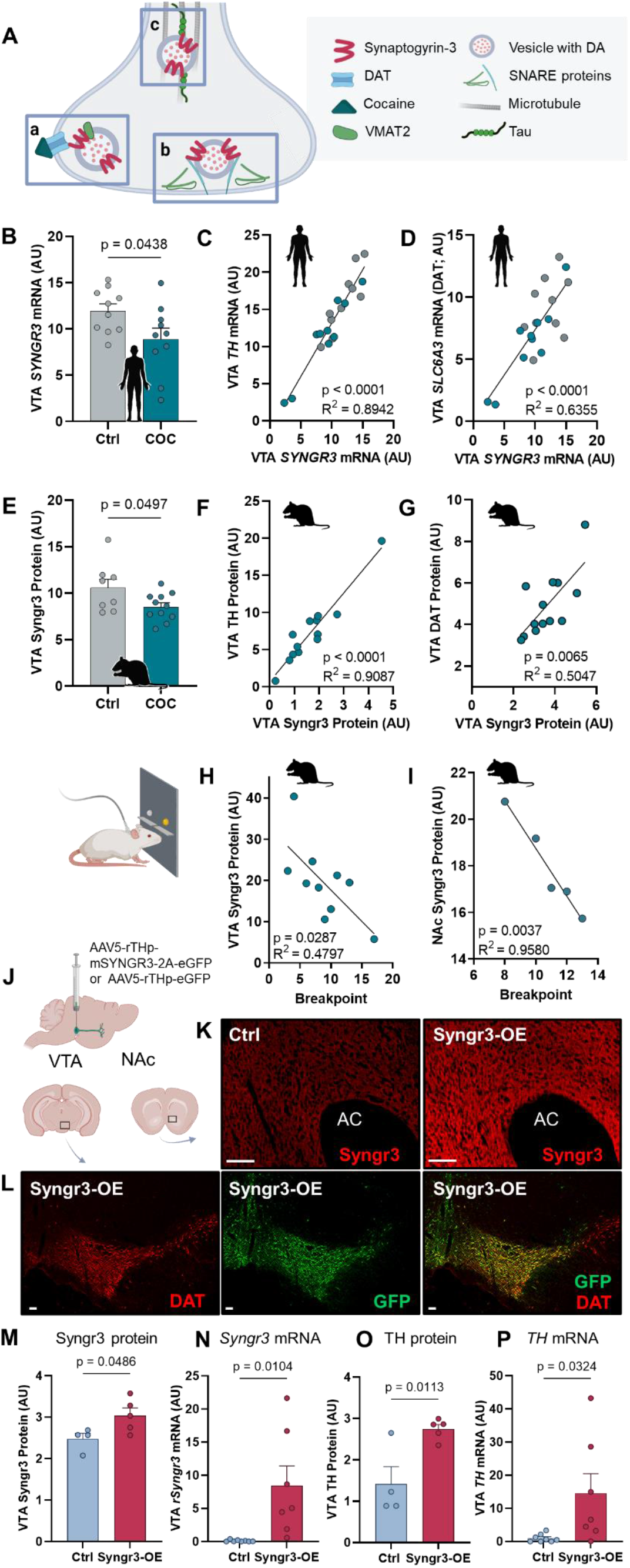
Interactions of mesolimbic Syngr3 levels with cocaine-directed behaviors and dopaminergic synaptic markers. (**A**) Schematic of Syngr3 localization in the presynaptic nerve terminal, where it interacts with (**Aa**) the DAT and vesicular monoamine transporter 2, (**Ab**) SNARE-related proteins, (**Ac**) and microtubule-associated tau. (**B**) *SYNGR3* mRNA expression was lower in postmortem VTA tissue from people with CUD. **C-D**, in both CUD (teal) and control (grey) groups, VTA *SYNGR3* mRNA positively correlated with (**C**) *TH* and (**D**) *SLC6A3* (DAT) mRNA. **E-G**, Similarly, rats that self-administered cocaine had (**E**) lower VTA Syngr3 protein levels, which positively correlated with (**F**) TH and (**G**) DAT VTA protein levels. **H-I**, in rats, Syngr3 protein levels negatively correlated with cocaine breakpoint in the (**H)** VTA and (**I**) NAc. (**J**) Schematic of Syngr3-OE viral infusion into VTA dopamine neurons. **K-L**, immunofluorescence of (**K**) Syngr3 (red) in NAc core terminals and (**L**) GFP (green) and the DAT (red) co-localization in the VTA (scale bar = 100 μm). **M-P**, VTA Syngr3 (**M**) protein and (**N**) mRNA levels are enhanced in Syngr3-OE rats. VTA TH (**O**) protein and (**P**) mRNA levels are also increased in Syngr3-OE rats. Error bars indicate mean ± SEM. VTA, ventral tegmental area; Syngr3, rat synaptogyrin-3 protein; *SYNGR3*, human synaptogyrin-3 mRNA; DA, dopamine; DAT, dopamine transporter; CUD, cocaine use disorder; COC, cocaine; SNARE, soluble N-ethylmaleimide sensitive factor attachment protein; VMAT2, vesicular monoamine transporter 2; TH, tyrosine hydroxylase 2; PR, progressive ratio; NAc, nucleus accumbens; Syngr3-OE, Synaptogyrin-3 overexpression; GFP, green florescent protein.

We found that postmortem VTA tissue from people who died of a cocaine overdose contained reduced levels of synaptogyrin-3 mRNA relative to healthy controls (*SYNGR3*; Fig. 1B; table S1 contains all statistical comparisons), and that *SYNGR3* levels positively correlated with the dopamine markers tyrosine hydroxylase (TH) and DAT (Fig. 1C-D). Similarly, rats that self-administered i.v. cocaine on an extended access paradigm^31^ had reduced VTA Syngr3 protein levels (Fig. 1E), which also correlated with TH (Fig. 1F-G). Moreover, we assessed motivation to self-administer cocaine by examining breakpoints on a progressive ratio (PR) reinforcement schedule^32^ and found that rats with the lowest mesolimbic Syngr3 levels had the highest motivation to seek and take cocaine (VTA: Fig. 1H; nucleus accumbens (NAc): Fig 1I). Thus, low Syngr3 levels were associated with two cardinal symptoms of CUD: high drug intake and high motivation to seek cocaine^33,34^.

To understand if low Syngr3 levels in dopamine neurons drove high cocaine breakpoints, we examined whether elevating Syngr3 expression selectively in VTA dopamine neurons (Syngr3-OE) would protect against the development of CUD-like behaviors (Fig. 1J). Syngr3 overexpression in the NAc (Fig. 1K) and VTA (Fig. 1L) was validated with immunofluorescence imaging. Additionally, Syngr3 protein (Fig. 1M) and mRNA (Fig. 1N) levels were significantly elevated in the VTA of Syngr3-OE rats. TH levels were also increased (Fig. 1O-P), suggesting that dopamine levels and activity may be enhanced as well.

Following acquisition of cocaine self-administration (Fig. 2A), Syngr3-OE rats displayed markedly reduced responding for cocaine under an unlimited access fixed ratio 1 (FR1) schedule (Fig. 2B). Not only did Syngr3-OE reduce cocaine intake per session, but it prevented escalation of intake over time (Fig. 2C), suggesting that Syngr3-OE blocked the progression to compulsive cocaine seeking and taking^33^. Syngr3-OE also decreased cocaine breakpoints on a PR schedule (Fig. 2D), indicating reduced motivation to work for cocaine^32,34^. Thus, augmented Syngr3 levels, specifically in dopamine neurons, attenuated CUD-like behaviors.

**Fig. 2.**
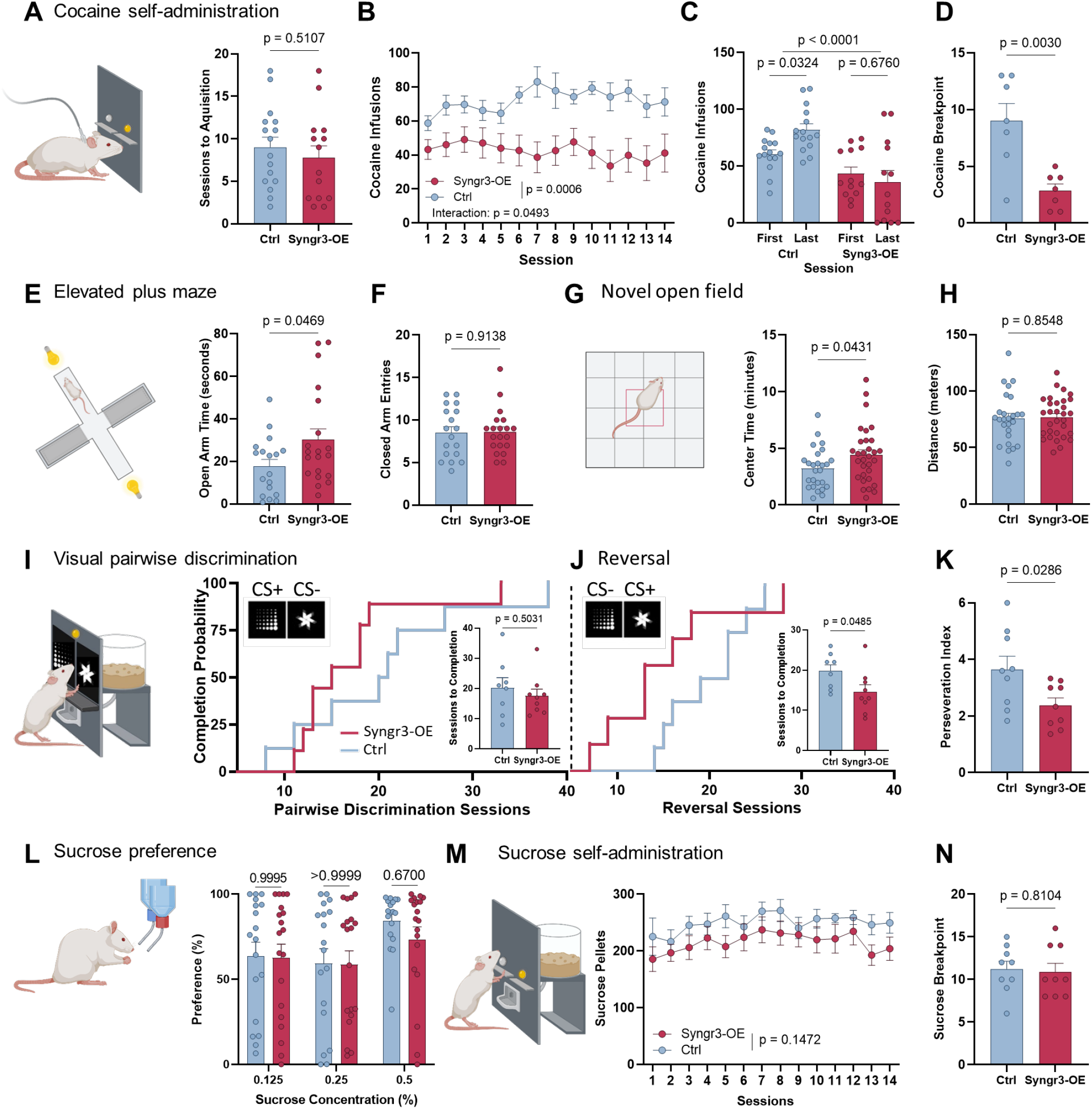
Synaptogyrin-3 overexpression (Syngr3-OE) specifically reduced cocaine self-administration and anxiety-like behaviors and enhanced cognitive flexibility. (**A**) Syngr3-OE rats acquired cocaine self-administration at the same rate (0.75 mg/kg/inf. IV cocaine, FR1, 6 hours/day, 20 infusion cap), (**B**) but took less cocaine on an unlimited access schedule of reinforcement (0.75 mg/kg/inf., FR1, 6 hours/day, unlimited). (**C**) Syngr3-OE prevented the escalation of cocaine intake and (**D**) decreased cocaine breakpoint (0.19 mg/kg/inf. cocaine, PR). Additionally, (**E**) Syngr3-OE rats spent more time on the open arms of an elevated plus maze, (**F**) but made the same number of closed arm entries. (**G**) Syngr3-OE rats also spent more time in the center of the novel open field, (**H**) but showed no difference in locomotion. **I-K**, On a pairwise discrimination task, (**I**) Syngr3-OE did not change the number of sessions to reach acquisition, but when the conditioned stimuli were reversed, (**J**) Syngr3-OE increased the speed of reversal learning by (**K**) reducing perseveration on the original conditioned stimulus. **L-M**, Syngr3-OE had no effect on (**L**) sucrose preference or responding on (**M**) unlimited access FR1 or (**N**) progressive ratio sucrose self-administration schedules. FR1, fixed ratio 1; PR, progressive ratio.

Anxiety disorders are often antecedent to and comorbid with CUD^35–37^. In preclinical models, high anxiety-like behavior is often associated with high cocaine intake^38–40^. To understand whether the low cocaine addiction-like behaviors documented in Syngr3-OE rats were accompanied by low anxiety-like behaviors, cocaine-naïve Syngr3-OE rats were tested on the elevated plus maze and novel open field. Indeed, Syngr3-OE increased open arm time on the elevated plus maze (Fig. 2E-F) and augmented center time in the novel open field (Fig. 2G) but had no impact on locomotion (Fig. 2H). Together, this indicated that Syngr3-OE reduced anxiety-like behavior, which may contribute to the diminished cocaine self-administration in Syngr3-OE rats.

Cocaine self-administration requires the ability to learn an operant behavioral task, so underlying cognitive impairments can affect these behaviors. To ascertain whether the reduction in cocaine self-administration in Syngr3-OE rats was influenced by impaired cognition, we employed a visual pairwise discrimination procedure^41^. Syngr3-OE rats showed no differences in acquisition or discrimination of the initial task (Fig. 2I), indicating normal associative learning, memory, and attention^41,42^. We then reversed the conditioned stimuli to assess cognitive flexibility, which is deficient in CUD and may perpetuate drug use^43,44^. Reversal learning requires inhibition of responding to the original conditioned stimulus as well as the learning of a new stimulus association (Fig. 2I-K). Syngr3-OE rats completed the reversal learning task more quickly than controls (Fig. 2J), exhibiting reduced perseveration of responding to the original cue, which indicates greater cognitive flexibility. Reduced cognitive flexibility is associated with habitual or compulsive responding and can perpetuate repetition of prior behaviors, thereby reducing the likelihood of stopping drug use behaviors ^43,44^. Therefore, enhanced cognitive flexibility likely contributes to the resilient phenotype of Syngr3-OE rats.

We next probed whether the marked reduction in reward responding produced by Syngr3-OE was specific for cocaine versus a natural reward using two sucrose self-administration protocols. In a two-bottle choice sucrose preference task, Syngr3-OE rats and controls showed similar preferences at each concentration (Fig. 2L). Further, in an operant sucrose self-administration task, there was no difference in responding for sucrose on either unlimited FR1 or PR schedules (Fig. 2M-N). Altogether, Syngr3-OE appears to selectively inhibit cocaine-related behaviors, a desirable outcome for a potential target for future CUD pharmacotherapeutics.

Due to the protective behavioral effects and elevated dopamine markers in Syngr3-OE rats, we hypothesized that Syngr3 strengthens dopamine neurotransmission. Microdialysis studies showed elevated basal extracellular levels in the NAc of freely behaving cocaine-naïve Syngr3-OE rats (Fig. 3A-B). In isolated NAc dopamine terminal fields in brain slices, fast scan cyclic voltammetry (FSCV) revealed augmented dopamine release in response to electrical stimulation (Fig. 3C-G) and greater calcium sensitivity (Fig. 3H), along with faster uptake (Fig. 3I). These results suggest a more active, responsive dopamine release profile along with heightened temporal and spatial control through increased DAT activity, that could result in rapid, discrete signaling events. Elevated dopamine activity in the NAc is consistent with the reduced cocaine self-administration, diminished anxiety-like behavior, and enhanced cognitive flexibility in Syngr3-OE rats.

**Fig. 3.**
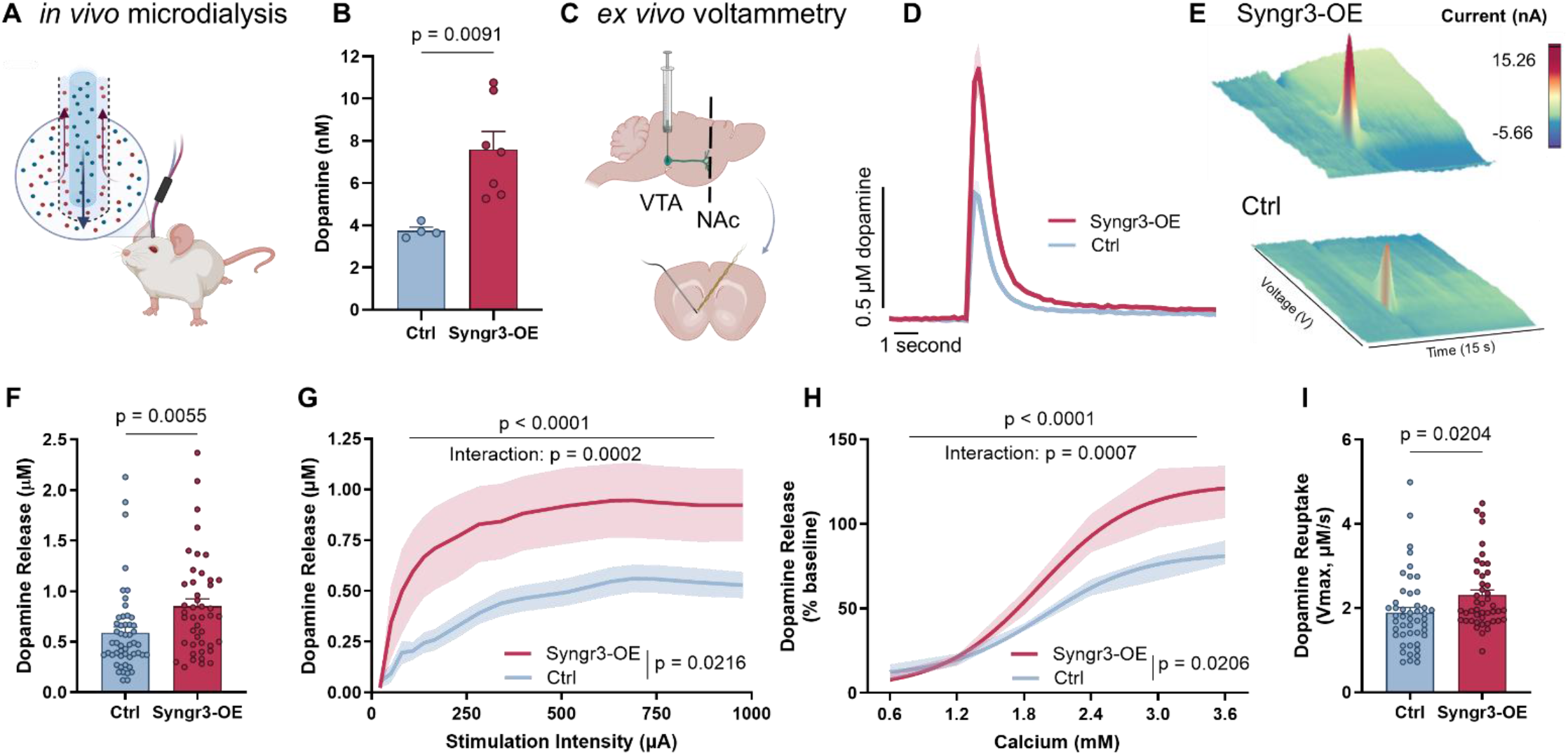
Synaptogyrin-3 enhanced NAc dopamine function. (**A**) Schematic showing *in vivo* microdialysis in cocaine-naïve Syngr3-OE and control rats, in which (**B**) basal dopamine levels were elevated in Syngr3-OE rats. (**C**) Schematic of *ex vivo* voltammetry in cocaine-naïve Syngr3-OE and control rats. **D-E**, Syngr3-OE rats had greater single-pulse stimulated dopamine release as shown with (**D**) concentration versus time 2D plot (averages of all baseline traces) and (**E**) a time (x-axis) versus voltage (y-axis) versus current (z-axis) plot (representative baseline traces). **F-G**, Syngr3-OE rats had increased dopamine release at (**F**) a perimaximal stimulation intensity (∼700 µA, same as **D**, see methods for stimulation details) and (**G)** across increasing applied currents. **H-I**, Additionally, (**H**) Release (at ∼700 µA) in Syngr3-OE rats was more sensitive to alteration in extracellular calcium concentrations and (**I**) had faster dopamine reuptake.

Decades of research have shown that chronic cocaine use induces a compensatory hypodopaminergic state that contributes to escalated drug use, greater drug seeking, and relapse-like behavior in humans and animals^5,6^. Since our work in cocaine-naïve Syngr3-OE rats suggested a more robust dopamine system, we postulated that Syngr3-OE may protect against cocaine-induced dopamine reductions. Therefore, we gave Syngr3-OE and viral control rats access to cocaine or saline in an extended access paradigm as above, then performed FSCV to assess NAc dopamine changes. Our group and others have previously shown that cocaine self-administration reliably reduces dopamine release^45,46^, which is observed in the viral control group here (Fig. 4B). However, virally augmented Syngr3 levels in mesolimbic dopamine neurons blocked this effect, as Syngr3-OE rats displayed similar dopamine release regardless of drug condition (Fig. 4C). Moreover, while cocaine reduced DAT function in the control group (Fig. 4D), there were no alterations in DAT function in the Syngr3-OE rats (Fig. 4E). Together, this demonstrates Syngr3’s ability to not only bolster dopamine function, but also to prevent cocaine-induced dopamine deficits.

**Fig. 4.**
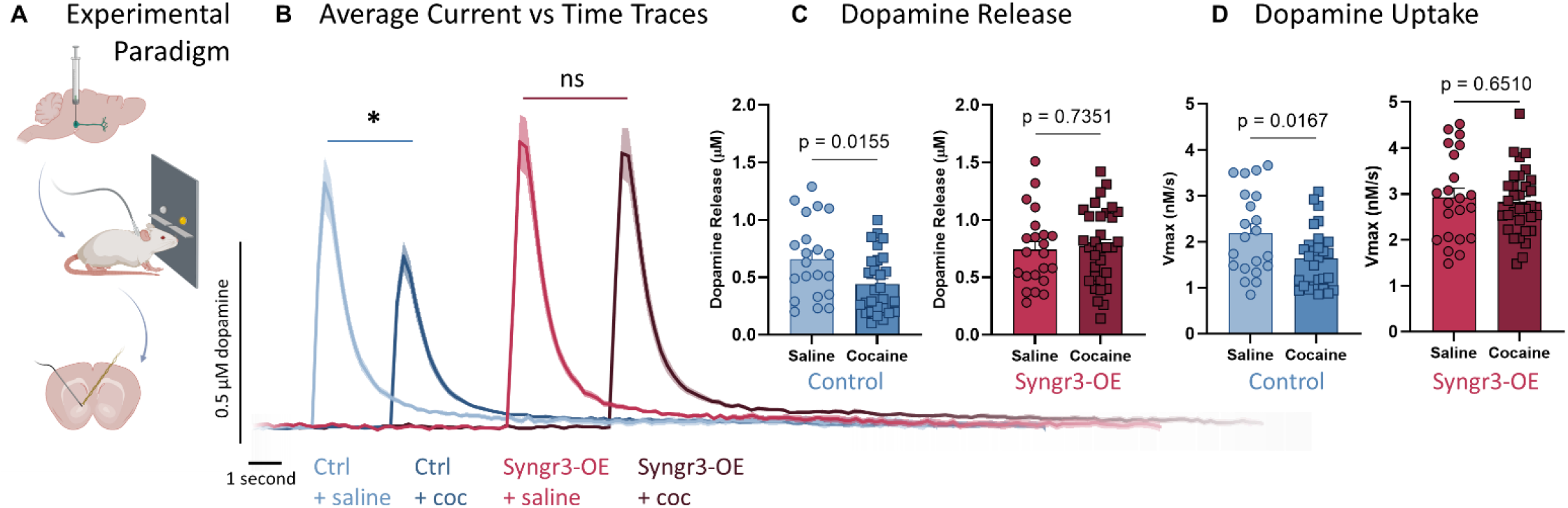
Synaptogyrin-3 prevented cocaine-induced dopamine deficits in the NAc. (**A**) Schematic showing viral infusion of Syngr3-OE or control virus, followed by cocaine or saline self-administration, then *ex vivo* voltammetry in the NAc core to measure dopamine kinetics. (**B**) Concentration versus time plots showing the average baseline traces of the four experimental groups: viral control rats that self-administered saline (light blue), viral control rats that self-administered cocaine (dark blue), Syngr3-OE rats that self-administered saline (light red), and Syngr3-OE rats that self-administered cocaine (dark red). (**C**) Cocaine self-administration reduced dopamine release among viral control rats (left) but had no effect on dopamine release in Syngr3-OE rats (right). (**D**) Similarly, cocaine self-administration slowed dopamine uptake in viral control rats (left), but not in Syngr3-OE rats (right). ** p < .01; ns, p > 0.05.

We postulate that Syngr3-OE acts by increasing the efficiency of dopamine neurotransmission, fine-tuning readily releasable pool dynamics, facilitating vesicular release and enhancing the recycling of released dopamine. This is consistent with Syngr3’s unique three-pronged influence in presynaptic terminals as, 1) an exocytotic vesicle protein^27,47^, 2) a DAT-associated protein^20,29^, and 3) a likely tau-mediated regulator of vesicle trafficking^21,28^ (Fig. 1A). This study reveals novel dopamine-enhancing and protective functions of the synaptic vesicle protein Syngr3 and provides mechanistic insight into its ability to prevent the progression to addiction-like behaviors in an animal model of CUD. Syngr3’s ability to bolster dopamine function without inducing hyperactivity or augmenting the rewarding effects of abused drugs makes it a potentially favorable target for development of pharmacotherapies to treat CUD. Additionally, in the small but growing number of publications that mention Syngr3, it has been linked to Parkinson’s disease^19,20,48^, Alzheimer’s disease^21,28^ and mild traumatic brain injury^22,49^, all of which involve dysregulated dopamine neurotransmission^50–54^, suggesting that this little-known protein may be important for normal dopamine function in a variety of anatomical locations and circumstances. Given the importance of proper dopamine signaling in multiple behaviors, including attention^55,56^, arousal^57^, motivation^58,59^, learning^60,61^, and motor planning^62,63^, discovering more about this novel protein may open important new avenues for therapies for a host of clinical indications.

## Materials and Methods

### Human postmortem brain tissue

We analyzed ventral tegmental area (VTA) synaptogyrin-3 (*SYNGR3*) mRNA expression from a publicly available dataset (GSE54839)^64^. The cocaine use disorder (CUD) cohort consisted of 10 people with a documented history of cocaine use and a positive postmortem toxicology report for cocaine (*i*.*e*., benzoylecgonine) at the time of death. The control cohort consisted of 10 age-matched subjects with no history of drug use who tested negative for cocaine and other drugs of abuse at the time of death.

The median expression from technical triplicates was normalized and analyzed using *limma* within the R programming language for statistical computing^65,66^ to determine differential gene expression (empirical bayes test) and inter-gene Pearson correlation coefficients.

### Animals

Male Sprague Dawley rats (325-350 g; Envigo, Indianapolis, IN) were pair-housed for at least one week while habituating in the vivarium. Rats were maintained on a reversed 12-hour light/dark cycle (lights on at 0300 hours) with food and water *ad libitum*, except for a subset of rats who were food restricted to 95% body weight for the duration of the pairwise discrimination procedure.

All animals were maintained according to the National Institutes of Health guidelines in Association for Assessment and Accreditation of Laboratory Animal Care accredited facilities, and experiments were performed in accordance with the Animal Care and Use Committee at Wake Forest University School of Medicine.

### Viral infusions

Rats were treated with meloxicam (1 mg/kg, subcutaneous (s.c.)) and isoflurane (induction: 4%; maintenance: 1.2-1.6%) for analgesia and anesthesia, respectively, then placed in a stereotaxic frame (Stoelting, Wood Dale, IL) equipped with StereoDrive (Neurostar, Tübingen, Germany). Rats received bilateral viral infusions into the VTA (AP: -5.6 mm, ML: ±0.7 mm, DV: -8.15 mm; 0° angle) to overexpress synaptogyrin-3 (Syngr3-OE; AAV5-rTHp-mSYNGR3-2A-eGFP, titer = 2.2 × 10^12^) or a viral control (elevated plus maze and novel open field experiments - rAAV5/TRUFR-eGFP CMV, titer = 2.2 × 10^12^; all other experiments - AAV5-rTHp-eGFP, titer = 3.7 × 10^12^). For all viral infusions, 0.6 µL virus was infused at 0.09 µL/minute, followed by a 10-minute wait for viral diffusion before removal of the injection needle. Rats received daily meloxicam (1 mg/kg, s.c.) for 48 hours after surgery and were individually housed for five weeks to allow viral expression. Following experiments, injection site placement was verified using histological visualization of green fluorescent protein (GFP) expression. All viral constructs were produced by the Vector Core of the Gene Therapy Center at the University of North Carolina (Chapel Hill, NC).

### Immunofluorescence

Rats were deeply anesthetized and transcardially perfused with phosphate-buffered saline (PBS; 0.12 M Na_2_HPO_4_, 0.18 M NaH_2_PO_4_, and 0.12 M NaCl; pH 7.4) followed by 10% formalin. Whole brains were removed and stored in 10% formalin at 4 °C overnight, then transferred to solutions of increasing sucrose concentrations (10-30% sucrose and 1% sodium azide in PBS; 4 °C) before being sliced into 35 µm thick coronal sections on a sliding microtome (American Optical Company, Buffalo, NY). Slices containing the nucleus accumbens (NAc) and VTA were incubated in a blocking solution containing PBS, 10% bovine serum albumin (BSA; Sigma-Aldrich Burlington, MA), and 0.5% Triton X-100 (Sigma-Aldrich). After two hours, primary antibodies for Syngr3 (Santa Cruz Biotechnology, Dallas, TX; sc-271046, 1:300), the dopamine transporter (DAT; Millipore Sigma, Burlington, MA; D6944, 1:250), and GFP (Aves Labs, Tigard, OR; AB2307313, 1:500) were added for an additional 16 hours at 4°C. The following day, slices were washed and incubated in a blocking solution containing Alexa 647 Goat-anti-Mouse (Abcam, Cambridge, UK, ab150115, 1:400), Alexa Flour Plus 555 Donkey-anti-Rabbit (Invitrogen, Waltham, MA; A32794; 1:400) and Alexa 488 Donkey-anti-Chicken (Jackson Immunoresearch, West Grove, PA; 703545155, 1:400) for two hours at room temperature. Finally, slices were mounted onto SuperFrost microscope slides and cover slipped using Aqua Poly/Mount (Polysciences, Warrington, PA). Slides were imaged on a Keyence fluorescence microscope (BZ-X710, Itasca, IL).

### Western blot hybridization

NAc and VTA tissue were rapidly dissected and frozen on dry ice. Total lysate was prepared by homogenizing tissue in T-PER (Thermo Scientific, Rockford, IL) and Halt Protease and Phosphatase Single Use Inhibitor Cocktail (Thermo Scientific), then agitating samples (one hour at 4 °C), and centrifuging (15,000 g for 20 minutes at 4°C). Protein concentrations were measured using a BCA protein assay kit (Thermo Scientific) and Molecular Devices Spectra Max 384 Plus spectrophotometer (Sunnyvale, CA) with SoftMax Pro software. Before loading, samples were heated for five minutes at 95°C, then 10μg of protein/sample and the Precision Plus Protein™ Kaleidoscope molecular weight ladder (BioRad, Hercules, CA) were added to 4-10% Mini-PROTEAN TGX gels (BioRad). Proteins from the gel were transferred to a nitrocellulose membrane (0.2 µm, BioRad), and bands were normalized using Revert 700 Total Protein Stain (Li-Cor Biosciences, Lincoln, NE). Membranes were washed and blocked with Intercept Blocking Buffer (Li-Cor Biosciences) for 45 minutes at room temperature. Then, primary antibodies for Syngr3 (Santacruz, sc-271046, 1:1000), the DAT (Millipore Sigma, D6944, 1:500), or tyrosine hydroxylase (TH; Millipore, AB152, 1:1000) were added to the blocking buffer and incubated for 16 hours at 4°C. The next day, membranes were treated with secondary antibodies, IRDye800CW Donkey-anti-Mouse (Li-Cor Biosciences, 926-32212; 1:400) and IRDye680RD Donkey-anti-Rabbit (Li-Cor Biosciences, 926-68073; 1:400), for two hours at room temperature. Membranes were imaged on an Odyssey Infrared Imaging System (Li-Cor Biosciences), and band analysis was done with Image Studio Lite (Li-Cor Biosciences).

The western blots in Figures 1H and 1I were run with slight modifications. Tissue was homogenized in T-PER and Protease Inhibitor Cocktail (Sigma-Aldrich, St. Louis, MO). The blocking buffer was 5% non-fat milk in TBST-0.1%, the primary antibody for Syngr3 was ab106460 (Abcam, 1:10,000), and the secondary antibody was Goat-anti-Mouse (Millipore Sigma, A8924, 1:1500). Digital images were captured with Chemi-Doc/Quantity One software (Bio-Rad), and Image Lab software (Bio-Rad) was used for analysis.

### RNA extraction for qPCR

For RNA extraction from the VTA, frozen tissue punches were briefly sonicated in Tris-buffered saline (TBS; 50mM Tris; pH 7.5). TBS homogenized tissue was lysed in RLT Lysis Buffer supplemented with ß-mercaptoethanol (Qiagen, Hilden, Germany). Subsequently, total RNA was isolated using an RNeasy plus Mini Kit according to the manufacturer’s instructions (Qiagen; cat. no. 74136).

### Real-time quantitative PCR (qRT-PCR)

Random-hexamer primed reverse transcription was performed using 1 µg total RNA per sample with the High-Capacity cDNA Synthesis kit (Applied Biosystems, Waltham, MA; cat. no. 4368814). cDNA was added to a reaction mix (5 µl total volume) containing 300 nM gene-specific primers and Power SYBR Green PCR mix (Applied Biosystems; cat. no. 4367659). All samples were run in triplicate and analyzed on a Quant Studio 5 PCR instrument (Applied Biosystems). Relative gene expression was calculated using the 2-^ΔΔCT^ method, with GAPDH gene expression serving as an internal control. Primer sequences (5’-3’) used are as follows: *Tyrosine hydroxylase (TH) F:* TGTCACGTCCCCAAGGTTCA; *TH R:* CTCTCCTCAAATACCACAGCCT; *Slc6a3 (DAT) F:* TGGCCATGGTGCCCATTTAT; *Slc6a3 R:* TGGCATAGGCCAGTTTCTCC; *Syngr3 F:* TTTGATCCCGTGAGCTTTGC; *Syngr3 R:* CGGCTATAGAAAACACCCAGG; Gapdh *F:* GTTTGTGATGGGTGTGAACC; *Gapdh R:* TCTTCTGAGTGGCAGTGATG. All primers were generated specifically for the rat genome.

### Elevated plus maze

Anxiety-like behavior was assessed in cocaine-naïve Syngr3-OE and control rats using a standard elevated plus maze (MED Associates, St. Albans, VT). The maze was raised 72.4 cm from the floor and consisted of two open runways with 1.3 cm high lips, and two closed runways with 40.6 cm high black walls. All runways measured 10.2 cm wide by 50.8 cm long and were connected by an open junction. A 7.5-watt light bulb was placed near the end of each open arm so that luminance measured 40 lux at the ends of the open arm and 2 lux in the junction.

After 60 minutes of habituation in the testing room, rats were placed in the central junction facing an open arm. Activity was monitored for five minutes via infrared beams and an overhead camera (Noldus, Wageningen, The Netherlands). All testing occurred during the second half of the dark (active) cycle (0900-1500 hours) under dim red light with white noise. Between each animal, the maze was cleaned, and luminance was checked and adjusted as needed.

### Novel open field

After one-hour habituation in the testing room, cocaine-naïve Syngr3-OE and control rats were placed in the center of individual chambers measuring 43 cm × 43 cm, with clear 30 cm high walls (MED Associates, St. Albans, VT) and monitored for 60 minutes. Locomotor activity and position in the field were measured with 16 × 16 photobeam arrays. The center was defined as 12 × 12 cm, and data from the first 15 minutes of testing were reported. Locomotion was defined as the total distance traveled. All open field testing occurred during the second half of the dark (active) cycle (0900-1500 hours) under dim red light and white noise. Chambers were thoroughly cleaned between each animal.

### Cocaine self-administration

#### Drugs

The National Institute on Drug Abuse Drug Supply Program generously provided Cocaine HCl (RTI, Log No.: 14201-142). Cocaine was dissolved in 0.9% sterile saline and filtered before experimental use.

#### Jugular vein catheterization

Rats were anesthetized with ketamine (100 mg/kg, intraperitoneal (i.p.)) and xylazine (10 mg/kg, i.p.) and treated with meloxicam (1 mg/kg, s.c.) for analgesia. A chronic silastic catheter was implanted in the right jugular vein, and the exit was mounted just caudal to the scapula as previously described^67^. Rats were individually housed in custom-made, operant chambers measuring 30 × 30 × 30 cm and received daily injections of meloxicam (1 mg/kg, s.c.) for 48 hours post-surgery. After three days of recovery, rats had their first self-administration session. All operant sessions occurred in the active/dark cycle (0900-1500), and catheters were flushed daily with heparinized saline (1.7 mg/kg, intravenous (i.v.)) to maintain patency. Cocaine-naïve control rats (Figure 1E) received catheters and underwent the same behavioral paradigms as cocaine self-administering rats but without the drug infusions; this controlled for surgery, tethering, housing, and operant cues (i.e., levers, cue lights, and pump sounds, etc.).

#### Acquisition/Training (all cocaine self-administering rats)

Rats had access to cocaine on a fixed ratio 1 schedule (FR1; 0.75 mg/kg/inf for six hours/day or until 20 infusions were reached). Immediately following each active response, the lever retracted for 20 seconds, a cue light above the active lever illuminated, and cocaine was intravenously infused at 0.025 mL/s for approximately four seconds (depending on body weight). Acquisition was defined as two consecutive days of 20 infusions.

#### Limited access cocaine self-administration (Figures 1 and 4)

Following acquisition, rats self-administered 1.5 mg/kg/inf cocaine under a limited access FR1 schedule for six hours per day or until 40 infusions were reached. As with training, a cue-light accompanied each active lever response, but the timeout was reduced to the length of the cocaine infusion (approximately four seconds). Once animals completed five consecutive days of 40 infusions, they were moved to the progressive ratio (PR) schedule of reinforcement.

Animals in Figure 4 underwent one additional day of limited access (40 infusions at 1.5 mg/kg/inf cocaine) after PR; they were sacrificed for voltammetry the following morning (∼18 hours after completion of the session).

#### Progressive Ratio (PR; all cocaine self-administering rats)

Animals completed either one day (Figure 1) or five days (Figure 2) on a PR schedule for 0.19 mg/kg/inf cocaine. Infusions were contingent upon progressively increasing response requirements over the course of the session: 1, 2, 4, 6, 9, 12, 15, 20, etc^32^. The outcome measure, breakpoint, was defined as the number of infusions obtained before one hour elapsed with no additional infusions earned or when the 6-hour session ended. If animals did not respond for cocaine within the first hour, the one-hour time limit started after the animals’ first lever press.

#### Unlimited access paradigm (Figure 2)

Syngr3-OE or viral control rats were placed on the acquisition procedure discussed previously (0.75mg/kg/inf cocaine for two consecutive days of 20 infusions). Once animals reached acquisition criteria, rats had unlimited access to cocaine (0.75mg/kg/inf) on an FR1 schedule for six hours per day. After 14 days of unlimited access, rats transitioned to the PR schedule.

### Pairwise Discrimination

#### Touchscreen Training

After sucrose self-administration (described below) and two weeks off in standard housing cages, cocaine-naïve Syngr3-OE and viral control rats were trained to touch stimuli presented on a computer touchscreen in operant chambers (Lafayette Instruments, Lafayette, IN). Each correct response resulted in the delivery of a 45 mg chocolate-flavored sucrose pellet (Bio-Serv, Flemington, NJ). Training occurred during the second half of their dark cycle (0900-1500) and involved four stages: initial touch training (ITT), must touch training (MTT), must initiate training (MIT), and punish incorrect training (PIT), as previously described^41^. Rats were required to complete one-hour sessions with a minimum of 30 trials and at least 80% accuracy for two consecutive days to move to the next stage. There were no differences in time to complete training sessions between SYG3-OE and control rats.

#### Pairwise Discrimination (PD) task

As previously described^41^, at the beginning of each one-hour session, a sucrose pellet was dispensed, and the magazine was illuminated. Upon retrieval of the sucrose pellet, the first trail was initiated, and two stimuli were presented on the digital screen: “fan” and “marbles”-shaped stimuli. One shape (either the “fan” or “marbles”) was assigned the sucrose-paired conditioned stimulus (CS+), and the other was paired with a timeout and house light (CS-). The shapes assigned CS+ and CS-were counterbalanced across rats, and the location of the shapes on the screen (right/left) were presented in a pseudorandom order, where the same locations were not presented for more than three consecutive trials. Touch on the CS+ resulted in a one-second tone and delivery of a sucrose pellet in the illuminated magazine, followed by a 10-second inter-trial interval (ITI) before the next trial could be initiated. Touch on the CS-terminated the trial, illuminated the house light for five seconds, and generated a 20-second ITI. After a CS-response, the subsequent trial(s) was a correction trial (CT), in which the stimuli were presented in the same location (right/left) until the CS+ was selected. CTs did not count towards the total number of trials but were used to calculate the perseveration index (the number of CTs divided by the number of incorrect responses on main trails^42^. Rats were considered to have acquired the PD task when they completed two consecutive days of at least 30 trials per session with 80% or more of their responses on the CS+.

#### Reversal of PD task

Following acquisition of the PD task, the CS+ and CS-were switched to assess cognitive flexibility. The sessions proceeded as described above. Rats were considered to have “reversed” after they performed two consecutive days of at least 30 trials per session with 80% or more of their responses on the new CS+.

### Sucrose Preference Task

Syngr3-OE or viral control animals without any previous behavioral testing were individually housed with access to two standard water bottles. On Monday, Wednesday, and Friday of the same week, water bottles were replaced with a bottle of sucrose (increasing concentrations: 0.125%, 0.25%, 0.5% in tap water) and a fresh bottle of tap water for two hours (1200-1400 hours). Bottle position alternated (right/left) across animals and days. A “drip cage” provided a measure of any sucrose or water loss during the placement of the bottles. Sucrose and water bottles were weighed before and after each session, and the drip cage value was subtracted from each rat’s total.

### Sucrose self-administration

Syngr3-OE and viral control animals without previous behavioral testing were individually housed in custom-made operant chambers that included a food hopper, magazine, and response levers. Animals were given access to sucrose pellets (FR1, 45 mg chocolate flavored sucrose pellets (Bio-Serv, Flemington, NJ), six hours/day, unlimited reinforced responses). Acquisition criteria were met after rats obtained at least 100 sucrose pellets and an active-to-inactive lever response ratio of at least 3:1 for two consecutive days. Following acquisition, rats were allowed to respond on an unlimited FR1 schedule for 14 days (FR1, 45 mg chocolate flavored sucrose pellets (Bio-Serv), six hours/day, unlimited reinforced responses). After the fourteenth session, rats were switched to a PR schedule for five days. Upon their last PR session, rats were returned to standard, non-operant cages for two weeks, then began pairwise discrimination (PD) training.

### *In vivo* microdialysis

Cocaine-naïve Syngr3-OE and viral control rats were anesthetized with ketamine (100 mg/kg, i.p.) and xylazine (10 mg/kg, i.p.) and given meloxicam (1 mg/kg, s.c.) for analgesia. Then, rats were head fixed in a stereotaxic apparatus (Stoelting), and a guide cannula (MD-2250; BASi Instruments, West Lafayette, IN) was implanted two mm above the NAc (AP: +1.6mm; L: +1.4mm; DV: -6.0mm; 0° angle). After surgery, concentric microdialysis probes (2 mm membrane, MD-2200; BASi Instruments, West Lafayette, IN) were inserted into the guide cannula. Rats were housed in testing chambers and recovered for approximately eight hours prior to starting “low-flow” microdialysis, during which a single sample was collected by continuously perfusing artificial cerebrospinal fluid (aCSF; NaCl 148 mM, KCl 2.7 mM, CaCl_2_ 1.2 mM, MgCl_2_ 0.85 mM; pH 7.4) at a flow rate of 0.5 µL/minute for 14 hours overnight (1900 to 1000 hours). The low flow rate allows the concentration of dopamine to reach equilibrium across the microdialysis membrane providing a sample of basal extracellular dopamine levels. Probe placement and viral placement/expression were validated via histology.

### High-performance liquid chromatography

Dopamine concentrations in dialysate were determined using high-performance liquid chromatography with electrochemical detection (HPLC-EC; ESA/Thermo Scientific, Chelmsford, MA). Separation of neurotransmitters and their metabolites was achieved using a reverse phase column (Luna 100 × 3.0 mm C18 3 μm, Phenomenex, Torrance, CA) with a mobile phase made up of 75 mM NaH_2_PO_4_, 1.7 mM 1-octanesulfonic acid sodium salt, 100 µL/L triethylamine, 25 µM EDTA, 10% acetonitrile v/v; pH 3.0. Analyte detection was carried out using a high-sensitivity analytical cell 5011 A (Thermo Fisher Scientific, Sunnyvale, CA) at +220 mV on a Coulochem III Electrochemical Detector (ESA/Thermo Scientific, Chelmsford, MA). Chromeleon software (ESA/Thermo Scientific, Chelmsford, MA) was used to quantify analytes using an external calibration curve of standards of known dopamine concentrations.

### *Ex vivo* fast scan cyclic voltammetry

Syngr3-OE or viral control rats were anesthetized with isoflurane, and the brain was rapidly removed and immersed in chilled, oxygenated aCSF containing (in mM): NaCl (126), KCl (2.5), NaH_2_PO_4_ (1.2), CaCl_2_ (2.4), MgCl_2_ (1.2), NaHCO_3_ (25), glucose (11), L-ascorbic acid (0.4). As previously described^69^, coronal brain slices (400 µm) containing the NAc core were collected using a vibrating tissue slicer (Leica VT1200S, Leica Biosystems, Wetzler, Germany), transferred to a recording chamber, and submerged in a bath of oxygenated aCSF (32 °C; perfused at a rate of 1 mL/minute). A bipolar stimulating electrode was placed in the NAc core, and dopamine was electrically evoked every three minutes (700 µA; four milliseconds; monophasic). A carbon-fiber microelectrode (CFE; 100–200 μM length, 7 μM diameter) was adjacent to the stimulating electrode, and a triangular waveform (−0.4 to +1.2 to −0.4 V vs. Ag/AgCl, 400 V/s) was applied to oxidize and reduce dopamine. Once dopamine signals were stable (at least 1 hour of baseline collections and <10% peak height variability), baseline dopamine release (µM) and uptake (µM/s) were determined, and the following experiments were run.

#### Input/output

After a baseline signal was established, the inter-stimulus interval was shortened to 60 seconds, and single pulse stimulations were applied at increasing voltage inputs (1.25, 1.5, 1.75, 2, 2.25, 2.5, 2.75, 3, 3.5, 4,5, 5, 6, 7, 7.5, 8, 9, 10 V) to optically isolated constant current output stimulations (Neurolog NL800A, Digitimer, Ft Lauderdale, FL, USA) calculated using the equation: Y = 0.0001158*X-0.0001808, where X=voltage input (V) and Y=current output (A), derived from the average calibration of all Neurologs used. Dopamine release (µM) was quantified at each stimulation intensity.

#### Calcium recording

Calcium sensitivity was assessed by bath applying progressively greater concentrations of Ca^2+^ and lesser Mg^2+^ to maintain extracellular divalent cation concentration (in mM: 0.6 Ca^2+^/ 3.0 Mg^2+^, 1.2 Ca^2+^/ 2.4 Mg^2+^, 1.8 Ca^2+^/ 1.8 Mg^2+^, 2.4 Ca^2+^/ 1.2 Mg^2+^, 3.0 Ca^2+^/ 0.6 Mg^2+^, 3.6 Ca^2+^/ 0.0 Mg^2+^).

#### Analysis

Electrodes were calibrated using a multiple linear regression model correlating the background charging current of each electrode with its sensitivity^70^. A unique set of regression coefficients were generated for the model from over 100 electrodes made within the lab and calibrated using the traditional *in vitro* method^70^.

Demon Voltammetry and Analysis software^71^ was used to analyze all FSCV data. Michaelis-Menten modeling was used to determine the concentration of dopamine released and the maximal rate of uptake (V_max_) following electrical stimulation.

### Statistical analysis

GraphPad Prism 9 (Graph Pad Software, La Jolla, CA) was used for all analyses and plots. Data are reported as mean ± SEM, and a significance was determined at *p* < 0.05. Specific statistical information, including tests used and sample size, is located in Supplementary Table 1. For molecular analyses (western blots, RT-qPCR), behavioral analysis (novel open field, EPM, operant self-administration, sucrose preference, pairwise discrimination), and microdialysis studies, all animals used were included as separate *N*s (i.e., samples were not pooled). For fast scan cyclic voltammetry, each slice represented an *n* (with approximately four slices per rat). Viral placement was confirmed in all rats, and none were excluded for missed targets. Data were excluded from the final analysis only if determined to be a significantly significant outlier (Grubbs test, *p* < 0.05) or for one of the following reasons:

*Cocaine self-administration:* animals were excluded if they failed to meet acquisition criteria (Syngr3: 4; Control: 2) or lost catheter patency (Syngr3: 5; Control: 4).

*Pairwise Discrimination:* one animal was excluded after failure to acquire the pairwise task (Syngr3: 0; Control: 1).

*In vivo microdialysis:* Sample contamination overlapping with dopamine detection parameters (Syngr3: 0; Control: 2).

*Western Blots:* not enough protein in sample (NAc sample from Figure 1I: 1)

---

**Table S1.**
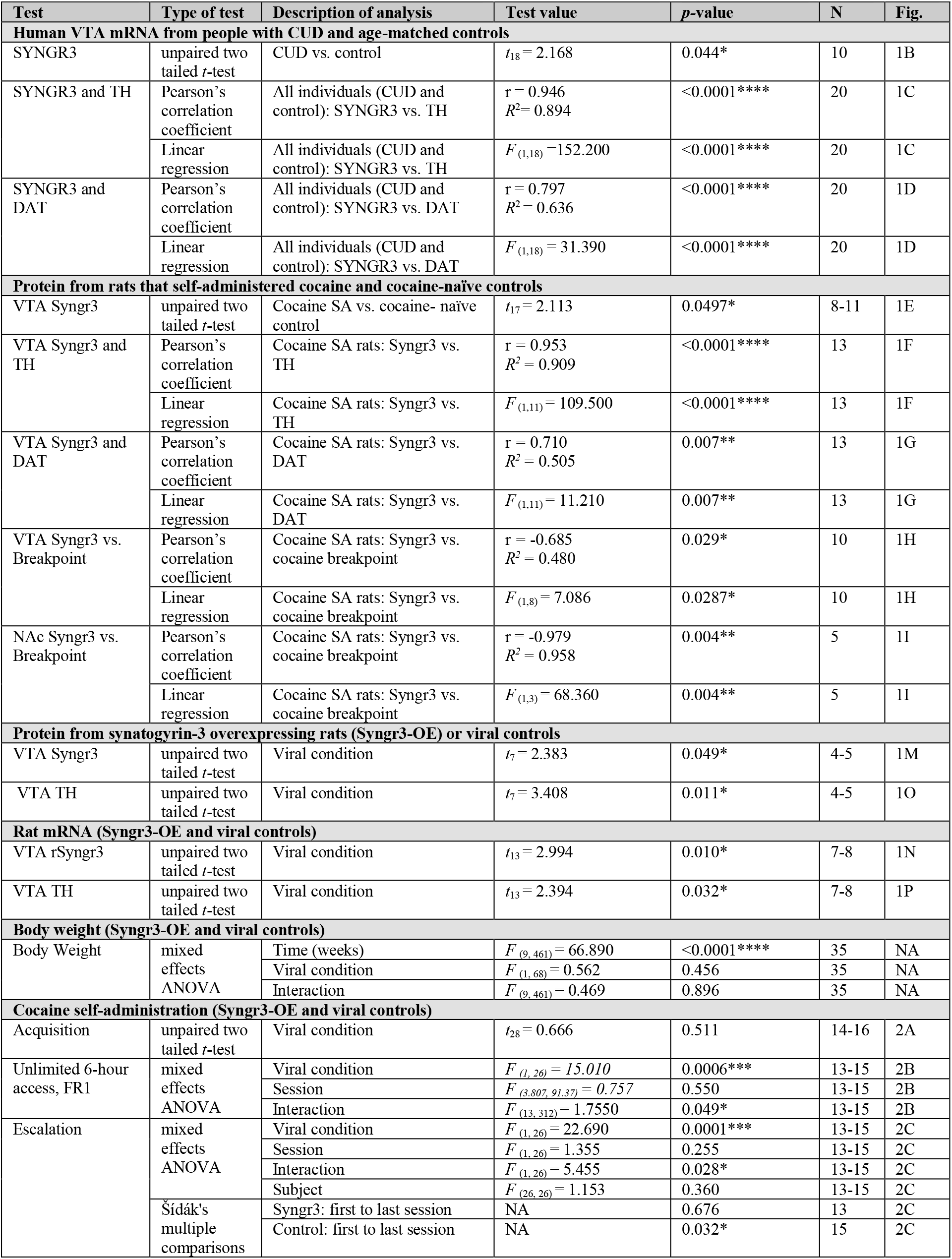
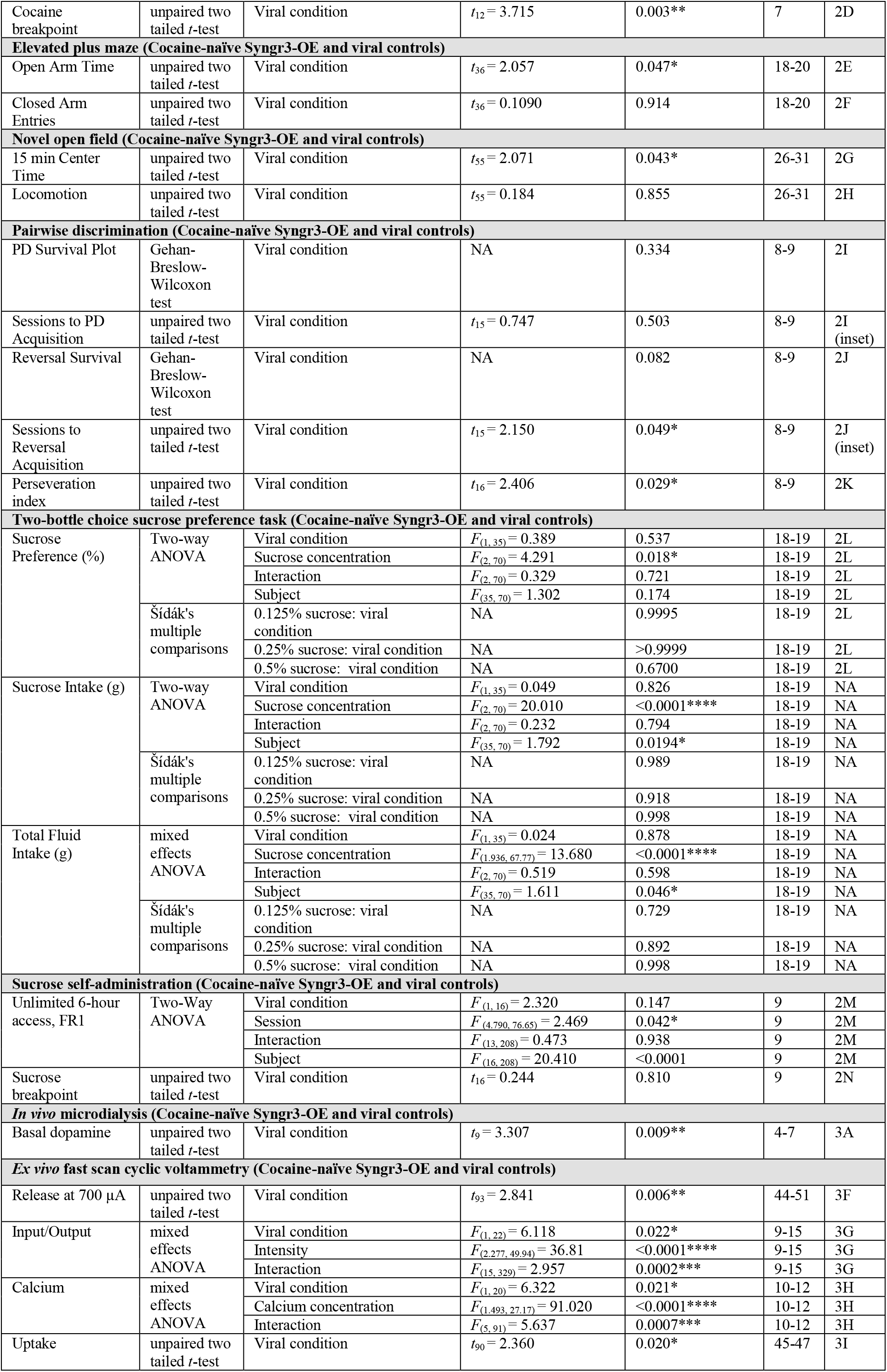
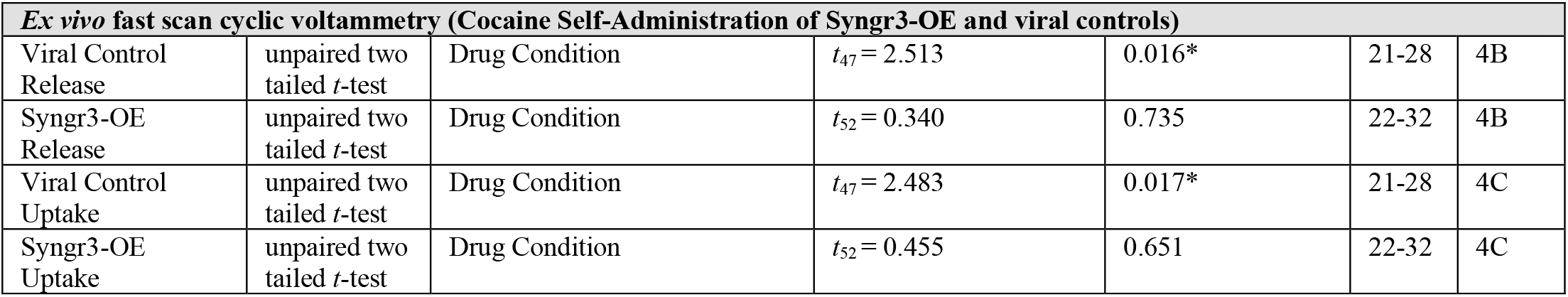
Statistical Results. All statistical results are listed in this table, grouped by experiment. Posthoc comparisons that failed to meet statistical significance with alpha set to 0.05 have been excluded from this table for brevity. VTA, ventral tegmental area; CUD, cocaine use disorder; SYNGR3, human synaptogyrin-3 gene; Syngr3, rat synaptogyrin-3 protein; TH, tyrosine hydroxylase; DAT, dopamine transporter; NAc, Nucleus accumbens; NA, not applicable; PD, pairwise discrimination; ANOVA, analysis of variance. * *p* < .05; ** *p* < .01; *** *p* < .005; **** *p* < .0001

## Acknowledgments

We thank Drs. Erin Calipari, Kim Raab-Graham, Jeff Weiner, Rebecca Hofford, Lindsey Kuiper, and Jordan Yorgason as well as Anna Neel and Samuel Barth for their support. Graphics were done with BioRender (full license to publish). This research was funded by the National Institutes of Health R01 DA054694 (to S.R.J.), P50 DA006634 (to S.R.J.), T32 DA041349 (to S.R.J., E.G.P., K.M.Holter, B.E.G., M.H.D.), F31 DA053105 (to E.G.P.), P50 AA026117 (to B.A.M.), R01 AA014445 (to B.A.M.), R01AA03067 (to B.A.M.), T32 AA007565 (to B.A.M, A.M.W.), R00 DA042129 (to R.W.G.), R00 AA024836 (to S.P.F.), U01 AA020889 (to S.P.F.), R01 AA030257 (to S.P.F.), T32 NS115704 (to O.A.O, A.M.C), and Fulbright Denmark (E.G.P).

## Author information

### Authors and affiliations

**Department of Physiology and Pharmacology, Wake Forest University School of Medicine, Winston-Salem, NC, USA**.

Emily G. Peck, Katherine M. Holleran, Kimberly M. Holter, Paige M. Estave, Jonathon P. Sens, Jason L. Locke, Olivia A. Ortelli, Brianna E. George, Monica H. Dawes, Alyssa M. West, Nancy J. Alexander, Drew D. Kiraly, Robert W. Gould, Brian A. McCool, Sara R. Jones

**Department of Psychiatry, Wake Forest University School of Medicine, Winston-Salem, NC, USA. ‘**

Drew D. Kiraly

**Department of Anesthesiology, University of Pittsburgh; Pittsburgh, PA, USA**.

Sean P. Farris

### Contributions

S.R.J., K.M.Holleran, and E.G.P. conceptualized the study. S.R.J., K.M.Holleran, and E.G.P. R.W.G, B.A.M, D.D.K., S.P.F. designed experiments. E.G.P., K.M.Holter, P.M.E., J.P.S., A.M.W., M.H.D., O.A.O., N.J.A., J.L.L., and B.M.E. collected data. E.G.P. made the figures and graphics. E.G.P., S.R.J., K.M.Holleran wrote the original draft. All authors contributed to review and editing.

### Corresponding author

correspondence to Sara R. Jones

## Ethics Declarations

### Competing interests

Authors declare no competing interests.

### Data and materials availability

The data presented in this study are available on request from the corresponding author.

## Notes

### Competing Interest Statement

The authors have declared no competing interest.

### Summary of Updates

Updated author information, funding, and figure references.

